# Decoding the Physical Mechanisms of Protein-Small Molecule Association by Multiscale Kinetic Monte-Carlo Simulations

**DOI:** 10.64898/2026.10.23.761358

**Authors:** Zhaoqian Su, Yinghao Wu

## Abstract

Protein-small molecule association involves multiple recognition stages governed by distinct physical interactions, yet resolving their kinetic contributions remains computationally challenging. Here, we develop a multiscale Monte Carlo framework that combines an atomistic, conformationally flexible ligand with a coarse-grained protein binding pocket and an atomistic steric envelope. Association is decomposed into three stages: molecular encounter, pocket entry, and productive binding. We evaluate the framework using ten protein–ligand complexes representing nine distinct targets, with experimentally derived association rates spanning nearly four orders of magnitude. Simulated pocket-entry rates exhibit the strongest correspondence with experimental kinetics (Pearson r = 0.617; Spearman ρ = 0.758), whereas productive-binding rates show closer agreement in absolute magnitude. Systematic perturbations of electrostatic interactions, hydrophobic attraction, hydrogen bonding, and ligand torsional flexibility reveal stage-dependent sensitivities. Initial encounter is comparatively insensitive to these perturbations, while hydrophobic attraction and conformational flexibility influence pocket entry. Productive binding exhibits additional sensitivity to local interactions, with excessive hydrogen-bond strengthening potentially hindering native-pose formation. Probability-derived kinetic-stage barriers and normalized mechanism descriptors further reveal heterogeneous recognition behavior across protein targets. Together, this framework provides a computationally efficient and interpretable approach for investigating protein-ligand association pathways and their physical determinants across diverse molecular systems.

## 1. Introduction

Molecular recognition between proteins and small molecules underlies numerous biological processes [1, 2] and forms the molecular basis of modern therapeutics [3, 4]. Through highly specific interactions with their protein targets, small molecules modulate enzyme catalysis [5], receptor activation [6], transcriptional regulation [7], intracellular signaling [8], and targeted protein degradation [9]. Understanding how proteins recognize small molecules is therefore fundamental for elucidating biological mechanisms and for the rational design of therapeutic agents. Binding kinetics has emerged as an important determinant of molecular recognition beyond static structures and equilibrium binding affinity. The association rate constant, *k_on_*, determines how rapidly a ligand finds and binds its target, whereas the dissociation rate constant, *k_off_*, determines the lifetime of the resulting complex. Together they define the equilibrium dissociation constant through *K_D_*= *k_off_*/ *k_on_*. While equilibrium affinity has traditionally been the primary objective of drug optimization, increasing evidence indicates that association kinetics strongly influences target engagement, selectivity, pharmacological response, and therapeutic efficacy [10]. Consequently, predicting protein–ligand association kinetics has become an important challenge in both mechanistic biology and drug discovery.

Despite this importance, computational studies of protein–ligand association kinetics remain far less developed than those of protein structure or binding affinity. Advances in structural biology and artificial intelligence have greatly improved the prediction of protein structures, ligand binding poses, and equilibrium affinities, supported by extensive resources such as PDBbind [11], BindingDB [12], and ChEMBL [13]. In contrast, experimentally measured association rates are relatively scarce, are rarely linked systematically to experimentally determined holo structures, and lack standardized benchmark datasets. Existing computational approaches, particularly Brownian dynamics [14–19], efficiently estimate diffusion-controlled association rates but generally rely on rigid molecular representations, atomistic treatment of entire proteins, and manually defined reaction criteria, limiting their ability to model flexible ligands and to scale across chemically diverse protein–ligand systems. On the other hand, all-atom molecular dynamics simulations [20–27] can, in principle, capture spontaneous protein-ligand association and provide estimates of association kinetics. Specialized high-performance platforms such as Anton [28, 29] have enabled direct observation of ligand-binding events in selected systems. However, these simulations typically require very long trajectories and substantial computational resources, making them impractical for routine application across large and chemically diverse protein-ligand datasets.

Here we present a multiscale kinetic Monte Carlo (KMC) framework for studying protein–small-molecule association. We first assembled a curated benchmark by integrating experimentally determined holo protein–ligand structures with experimentally measured dissociation rates and equilibrium binding affinities, from which association rates were derived. We then developed an adaptive-resolution molecular representation in which only the protein binding pocket is modeled using a coarse-grained description, whereas the ligand is represented explicitly by all heavy atoms. Ligand flexibility is incorporated through torsional Monte Carlo sampling, and intermolecular interactions are described using steric exclusion, van der Waals interactions, electrostatics, hydrophobic interactions, and hydrogen bonding. This adaptive representation substantially reduces computational cost while preserving the molecular detail required for ligand recognition.

Unlike conventional approaches that treat association as a single reaction event, our framework explicitly decomposes ligand recognition into three sequential kinetic stages: diffusional encounter, pocket entry, and productive binding. The model estimates both the rate of each stage and the transition probabilities between successive stages, enabling the overall association process to be resolved into distinct kinetic barriers. More importantly, we extend the framework beyond rate prediction by using it as a computational perturbation platform. Individual physical interactions, including electrostatic interactions, hydrophobic interactions, hydrogen bonding, and ligand flexibility, are systematically removed, weakened, or strengthened while all other simulation conditions remain unchanged. The framework therefore generates experimentally testable hypotheses regarding protein conformational gating, solvent reorganization, kinetic trapping, or other mechanisms that cannot be inferred from equilibrium structures alone. Together, this work establishes an interpretable multiscale framework for studying protein–ligand association kinetics by integrating a curated benchmark, adaptive-resolution KMC simulation, multistage kinetic decomposition, and systematic energy perturbation analysis. Beyond predicting association rates, it provides a quantitative platform for decoding the physical mechanisms of protein–ligand recognition and lays the foundation for future physics-informed machine learning, kinetic virtual screening, and mechanistic optimization of therapeutic ligands.

## 2. Model and Methods

### 2.1. Benchmark Dataset Construction

We constructed a benchmark of protein–ligand association kinetics by integrating kinetic, affinity, and structural data from multiple public resources. Because experimentally measured *k_on_*values are rarely available together with corresponding holo structures, the association rates used in this benchmark were reconstructed from independently reported *k_off_* and *K_D_*measurements. Consequently, some complexes may reflect differences in experimental conditions or assay protocols, and the benchmark is intended primarily for evaluating relative kinetic trends and mechanistic behavior rather than absolute quantitative agreement. Specifically, experimentally measured dissociation rate constants (*k_off_*) were obtained from the curated dataset reported by Liu et al. [30]. Corresponding equilibrium dissociation constants (*K_D_*) were retrieved from BindingDB [12] by matching protein targets (UniProt identifiers) and ligand identities. Association rate constants were then estimated as *k_on_* = *k_off_* / *K_D_*. Candidate complexes were subsequently mapped to experimentally determined holo structures in the Protein Data Bank (PDB). Only complexes satisfying all of the following criteria were retained: (i) an experimentally resolved holo structure containing the bound ligand, (ii) consistent protein and ligand identities between the kinetic and structural data, (iii) complete structural information suitable for KMC simulations, and (iv) unambiguous ligand connectivity. Complexes with missing coordinates, incompatible ligands, ambiguous mappings, or multicomponent ligand representations were excluded. The final benchmark comprises a curated collection of experimentally characterized protein–ligand complexes spanning multiple protein families and ligand chemotypes, each paired with a holo structure and an experimentally derived association rate for evaluating KMC simulations.

### 2.2. Adaptive-Resolution Molecular Representation

Each protein-ligand system in the dataset was constructed directly from its experimentally determined holo structure. The native ligand was identified by matching the chemical identity of the ligand in the kinetic dataset to the corresponding PDB Chemical Component Dictionary entry.

Full stereochemical InChIKey matching was used whenever possible; when the input ligand lacked complete stereochemical specification, matching was performed using the connectivity block of the InChIKey, while retaining the experimentally resolved ligand coordinates from the holo structure.

To balance computational efficiency with the chemical resolution required for small-molecule recognition, the protein and ligand were represented at different levels of detail (left panel of **Figure 1**). The ligand was represented explicitly by all non-hydrogen atoms. Its covalent connectivity and bond topology were obtained from the PDB Chemical Component Dictionary, and the idealized coordinates were replaced by the experimentally observed coordinates in the holo structure. Partial atomic charges were assigned using the Gasteiger procedure. Hydrogen-bond donor, hydrogen-bond acceptor, and hydrophobic atom identities were assigned using RDKit chemical-feature definitions. Rotatable bonds were identified as non-ring single bonds satisfying a standard rotatable-bond substructure definition.

**Figure 1.**
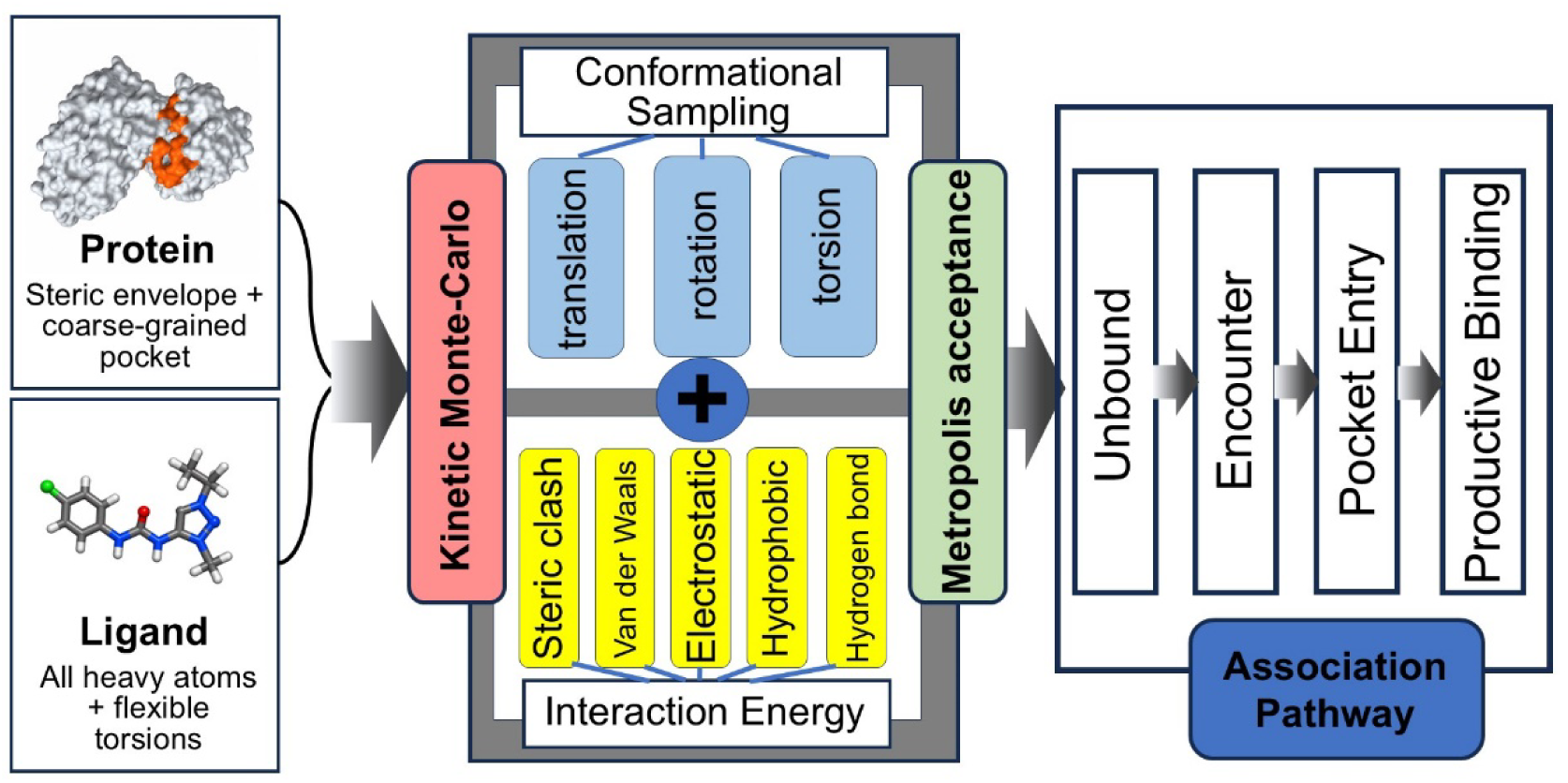
Schematic overview of the multiscale Monte Carlo framework for protein–small-molecule association. The framework combines a flexible ligand represented by all heavy atoms with a coarse-grained protein binding pocket and an atomistic protein steric envelope. Ligand configurations are sampled through translational, rotational, and internal torsional Monte Carlo moves, with acceptance determined by an effective interaction potential incorporating steric exclusion, van der Waals interactions, electrostatics, hydrophobic attraction, and hydrogen bonding. Molecular association is resolved into three sequential recognition stages: initial encounter, pocket entry, and productive binding. Stage-specific apparent association rates and conditional transition probabilities are estimated from independent simulation trajectories.

The protein binding pocket was represented using a residue-level coarse-grained description. Pocket residues were defined as residues containing at least one heavy atom within 7.0 Å of any ligand heavy atom in the native holo structure. Each selected residue was represented by up to two interaction sites. A backbone site was placed at the residue Cα position, and a side-chain site was placed at the geometric center of the side-chain heavy atoms. The backbone site was assigned a neutral charge and weakly hydrophilic character, whereas the side-chain site carried residue-specific charge, hydrophobicity, and hydrogen-bond donor or acceptor annotations. Aspartate and glutamate were assigned negative charges, lysine and arginine positive charges, and histidine a weak positive charge. Residue hydrophobicities were represented by an empirical scale ranging from strongly hydrophobic aliphatic and aromatic residues to charged hydrophilic residues.

Although energetic interactions were evaluated primarily between ligand atoms and coarse-grained pocket sites, all protein heavy atoms were retained in a spatial search tree for global steric exclusion. This hybrid representation prevented the ligand from penetrating non-pocket regions of the protein while avoiding the computational expense of evaluating detailed chemical interactions against every protein atom. The resulting model therefore combined an all-heavy-atom flexible ligand, a chemically annotated coarse-grained pocket, and an atomistic protein envelope for excluded-volume enforcement.

The experimentally observed ligand pose was used to define the native reference state. A ligand atom-pocket site pair was designated as a native contact when its separation in the holo structure was no greater than 4.5 Å. These native contacts were used only to characterize progression toward productive binding and were not introduced as an explicit structure-based attractive bias in the interaction potential.

### 2.3. Effective Interaction Energyy

The interaction energy between a protein and its ligand with configuration X was represented as the sum of five physically interpretable terms (middle panel of **Figure 1**):

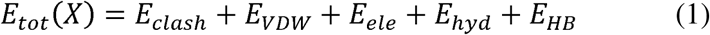

This potential was designed as an efficient effective interaction model rather than a complete all-atom molecular-mechanics force field. It preserves the dominant interactions expected to guide ligand diffusion, pocket capture, and native-pose stabilization while allowing large numbers of independent association trajectories to be generated. In specific, the first term on the right hand side of Equation (1) accounts for the global protein-ligand steric overlap. It was evaluated using the distance between each ligand heavy atom *i* and its nearest protein heavy atom:

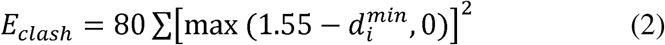

This soft quadratic penalty is zero when the nearest protein atom lies beyond 1.55 Å and rises rapidly at shorter separations, preventing unphysical penetration of the protein interior.

The second term describes the short-range packing between ligand atoms and pocket sites using a softened Lennard-Jones-like potential:

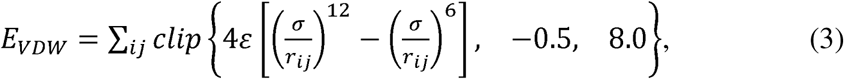

where *r_ij_*is the distance between ligand atom *i* and pocket site *j*, σ = 3.4 Å, and ε = 0.08 kcal mol ¹. Distances were bounded below at 1.2 Å for numerical stability. Clipping the pairwise energy limited extreme repulsive values while retaining favorable short-range packing.

Electrostatic interactions were represented using a Debye-Hückel potential in the third term:

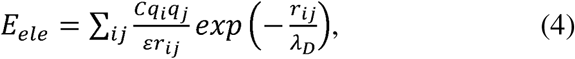

where *q_i_* is the Gasteiger charge of ligand atom *i*, *q_j_* is the assigned charge of pocket site *j*, C = 332.06371 converts the electrostatic energy to kcal mol ¹, and ε = 78.5 is the dielectric constant of water. The Debye screening length was calculated as

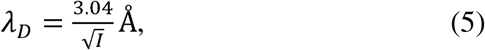

where *I* is the ionic strength in molar units. Simulations were performed at *I* = 0.15 M unless otherwise stated.

Hydrophobic effect of the protein-ligand interactions was captured by a smooth distance-dependent attraction in the fourth term:

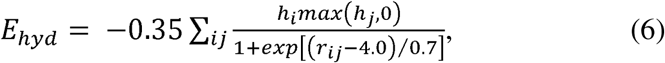

Here, *h_i_* is a binary indicator specifying whether ligand heavy atom *i* is classified as hydrophobic (*h_j_* =1) or nonhydrophobic (*h_j_* =0), based on *RDKit* chemical-feature definitions. In contrast, *h_j_* is a residue-dependent hydrophobicity parameter assigned to protein pocket site *j* according to the amino-acid identity represented by that site. Positive values denote hydrophobic pocket environments, whereas polar or charged residues have smaller or negative values. Only favorable hydrophobic contributions were included through max(*h_j_*, 0); thus, negative residue hydrophobicities were set to zero rather than treated as explicit repulsive interactions. The sigmoidal distance function smoothly switches the interaction on as a hydrophobic ligand atom approaches a hydrophobic pocket site, reaching half of its maximum magnitude at *r_ij_* =4.0Å and decaying toward zero at longer distances. This term therefore provides an effective representation of short-range hydrophobic capture and burial rather than an explicit solvent-mediated hydrophobic free energy.

Finally, hydrogen-bond interactions were represented by a distance-dependent donor-acceptor potential:

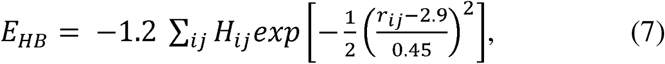

where *H_ij_* = 1 when ligand atom *i* and pocket site *j* form a complementary donor-acceptor pair and *H_ij_* = 0 otherwise. Ligand heavy atoms were classified as hydrogen-bond donors and/or acceptors according to RDKit chemical-feature definitions, whereas protein pocket sites were assigned donor and acceptor character according to the chemical properties of the corresponding amino-acid residue. The Gaussian potential is centered at 2.9 Å, corresponding to a favorable heavy-atom hydrogen-bond distance. The current implementation does not explicitly include hydrogen-bond angular dependence and should therefore be interpreted as an effective local stabilization term.

### 2.4. Kinetic Monte Carlo Simulation Algorithm

Independent KMC trajectories were initiated from unbound ligand configurations. The native ligand coordinates were first translated to their center of mass and randomly rotated. The ligand was then placed at a radial distance of 28 Å from the geometric center of the protein in a randomly selected direction. This procedure generated different initial positions and orientations while preserving the internal ligand conformation derived from the holo structure. The protein remained fixed throughout each trajectory.

At every simulation step, one of three ligand moves was selected: translation with probability 0.60, rigid-body rotation with probability 0.27, or internal torsion with probability 0.13. These probabilities were empirically chosen to emphasize translational diffusion during long-range association while maintaining substantial sampling of ligand orientation and internal conformation. For translational moves, a three-dimensional displacement vector was sampled from a normal distribution with a standard deviation of 0.60 Å in each Cartesian direction. For rotational moves, the ligand was rotated around its center of mass about a randomly selected axis, with the rotation angle sampled from a normal distribution with a standard deviation of 10°. For torsional moves, one rotatable ligand bond was selected randomly. The smaller of the two molecular fragments separated by that bond was rotated around the bond axis, with the torsional angle sampled from a normal distribution with a standard deviation of 22°. Rotations involving ring bonds were excluded. The proposal amplitudes were selected to permit efficient exploration while avoiding excessively large moves that would frequently generate steric clashes or low acceptance probabilities. Overall, this procedure allowed the ligand to adapt its internal conformation while preserving its bond lengths, bond angles, and molecular connectivity.

The ligand was confined within a spherical simulation region centered on the protein. If a proposed ligand center of mass crossed the outer radius of 38 Å, the displacement was reflected back into the simulation region. This reflecting boundary prevented irreversible ligand escape while avoiding periodic copies of the protein.

Following each proposed move, the interaction energy of the trial configuration expressed in Equation (1) was evaluated. A move with energy change Δ*E = E_trail_ - E_current_* was accepted according to the Metropolis criterion.

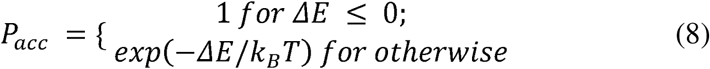

At temperature *T*, the thermal energy was calculated as *kT* = 0.0019872 *T* kcal mol ¹. For the baseline simulations, we uses *T* = 298.15 K.

Accepted configurations replaced the current configuration, whereas rejected moves left the ligand unchanged. Because each trajectory contained stochastic translations, rotations, torsional moves, and acceptance decisions, multiple independent trajectories were generated for every protein-ligand complex.

Following the simulations, we define different kinetic states along the association process (right panel of **Figure 1**). In current study, the association process was decomposed into three sequential kinetic states.

A trajectory reached the *encounter state* when the minimum distance between any ligand heavy atom and any pocket site was no greater than 6.0 Å (min*_ij_ r_ij ≤_* 6.0Å). The *pocket-entry state* was reached when the ligand center of mass was within 7.0 Å of the center of mass of the native ligand pose 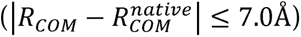. Finally, the productive-binding state required pocket entry together with both geometric and contact-based recovery of the native pose. Specifically, the ligand heavy-atom root-mean-square deviation from the native ligand coordinates was required to be no greater than 3.0 Å, and at least 40% of the native ligand-pocket contacts had to be recovered (*RMSD; ≤ 3.0*Å *Q_native_≥* 0.40Å).

To suppress transient fluctuations, the productive-binding condition had to remain satisfied for five consecutive KMC steps before a trajectory was classified as successfully bound. The first steps at which the encounter, pocket-entry, and productive-binding criteria were met were recorded as their respective first-passage steps. A trajectory was terminated after persistent productive binding or after reaching the predefined maximum number of simulation steps.

An approximate physical time increment was assigned to each KMC step from the translational diffusion relation:

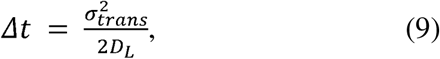

where σ*_trans_* = 0.60 Å is the translational proposal scale and *D_L_* = 50 Å² ns ¹ is the assumed ligand translational diffusion coefficient. This mapping provides an effective simulation clock for comparing first-passage behavior among systems. It should not be interpreted as an exact dynamical integration because rotational moves, torsional moves, rejected proposals, solvent friction, and protein conformational dynamics are represented only approximately.

For each kinetic endpoint x, where x denotes encounter, pocket entry, or productive binding, the event probability was calculated from successful trajectories *N_x_* among independent trajectories *N_tot_*:

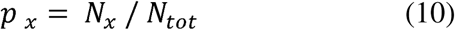

The apparent second-order rate constant was then estimated as

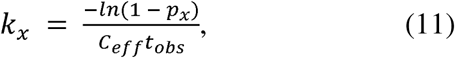

where *t_obs_*= *N_step_*×Δ*t* is the maximum observation time and *C_eff_*is the effective single-ligand concentration associated with the spherical initial volume:

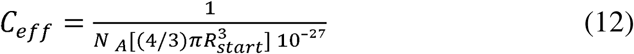

with *R_start_* = 28 Å denoting the initial radial distance at which the ligand is placed from the protein center when each simulation trajectory begins and *N_A_*denoting Avogadro’s constant. This procedure yielded three apparent rates, *k_encounter_*, *k_entry_*, and *k_productive_*, which characterize successive stages of molecular recognition rather than treating association as a single event.

### 2.5. Code availability

All relevant source codes can be found in the GitHub repository: https://github.com/wujah/PLassKMC/.

## 3. Results

### 3.1. A structurally and kinetically diverse protein–ligand benchmark

Independent Following the benchmark construction procedure described in the Methods section, we initially identified 11 protein–ligand complexes with matched experimental kinetic information and holo structures. One complex (PDB: 5A3O) was excluded because its ligand could not be unambiguously reconstructed as a single connected molecule for simulation. The final benchmark therefore comprised 10 protein–ligand complexes representing nine distinct protein targets (**Table 1** and **Figure 2**).

**Figure 2.**
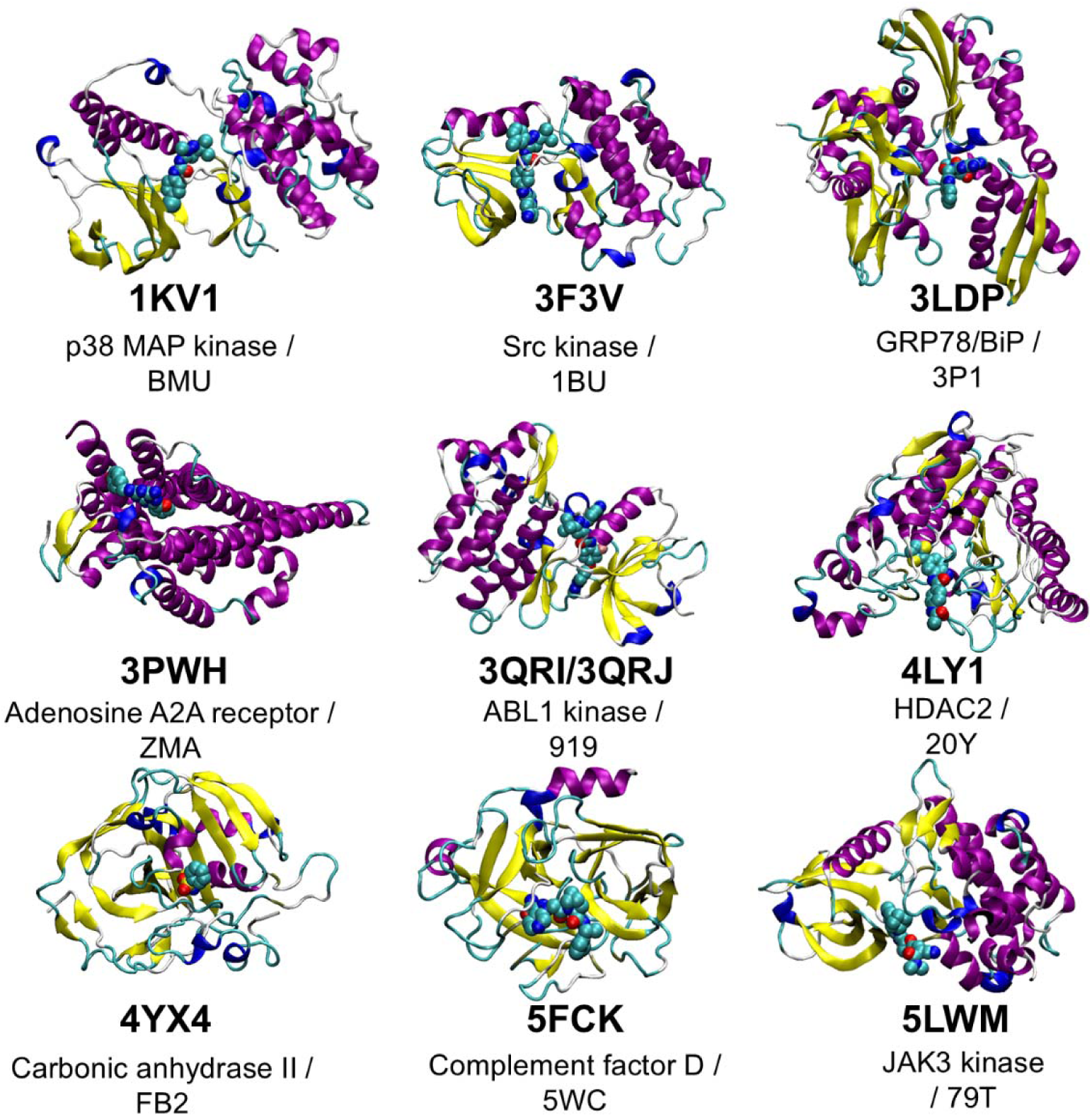
Structural diversity of the protein–ligand complexes included in the benchmark. Crystal structures of the ten protein–ligand complexes representing nine distinct protein targets. Proteins are shown in cartoon representation, with α-helices colored magenta, β-sheets yellow, and loops and connecting regions in cyan, blue, and gray. Bound ligands are displayed as space-filling spheres, highlighting their positions within the protein binding pockets. Each structure is labeled with its PDB identifier, protein target, and ligand chemical component identifier. The ABL1 wild-type and T315I mutant complexes are grouped together because they contain the same ligand. The structures illustrate the diversity of protein folds, ligand-binding environments, and molecular architectures represented in the benchmark.

**Table 1.**
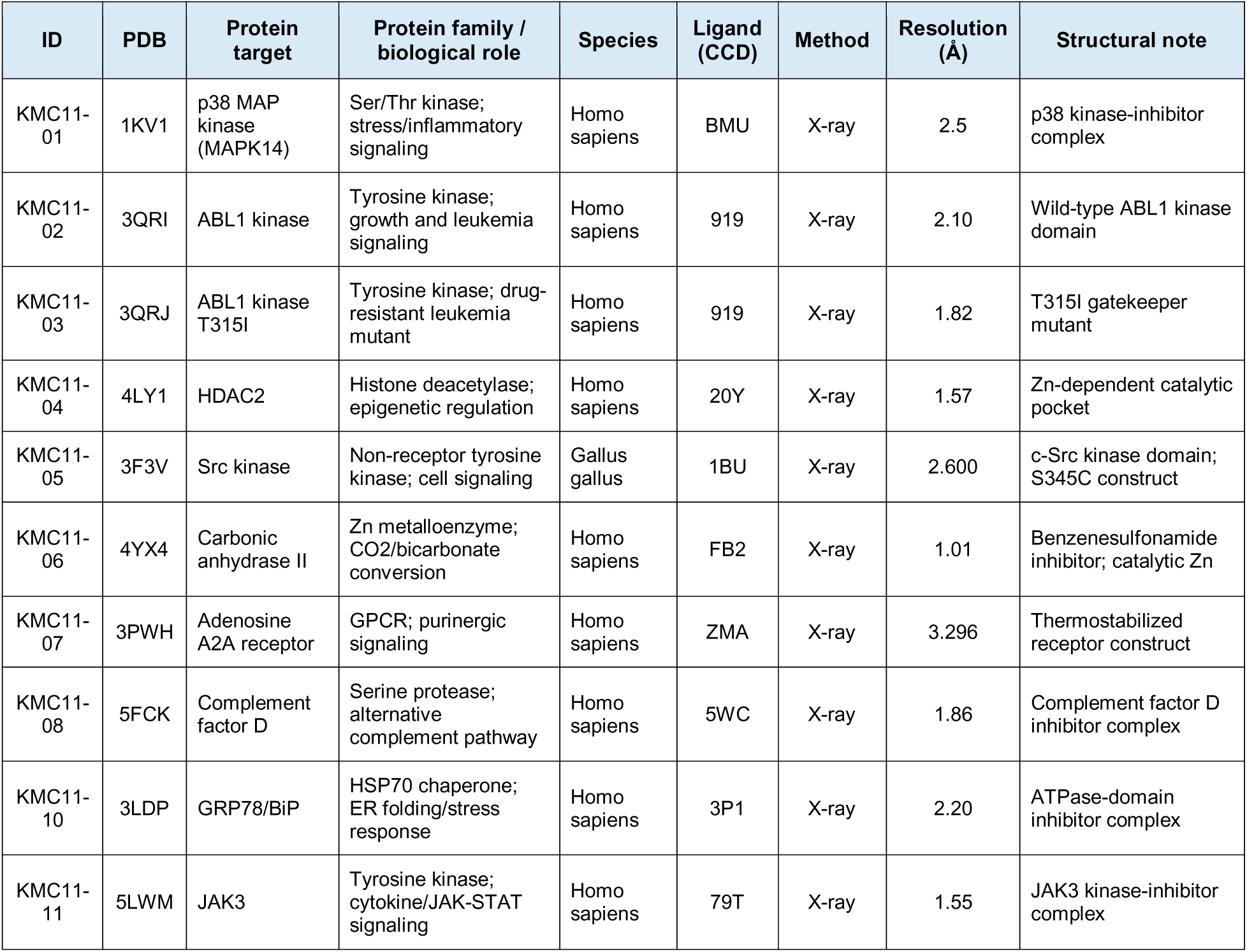
Structural characteristics of the protein–ligand benchmark. Summary of the ten protein–ligand complexes, including protein targets, biological functions, ligand identities, crystallographic resolutions, and structural features.

| ID | PDB | Protein target | Protein family / biological role | Species | Ligand (CCD) | Method | Resolution (Å) | Structural note |
| --- | --- | --- | --- | --- | --- | --- | --- | --- |
| KMC11-01 | 1KV1 | p38 MAP kinase (MAPK14) | Ser/Thr kinase; stress/inflammatory signaling | Homo sapiens | BMU | X-ray | 2.5 | p38 kinase-inhibitor complex |
| KMC11-02 | 3QRI | ABL1 kinase | Tyrosine kinase; growth and leukemia signaling | Homo sapiens | 919 | X-ray | 2.10 | Wild-type ABL1 kinase domain |
| KMC11-03 | 3QRJ | ABL1 kinase T315I | Tyrosine kinase; drug-resistant leukemia mutant | Homo sapiens | 919 | X-ray | 1.82 | T315I gatekeeper mutant |
| KMC11-04 | 4LY1 | HDAC2 | Histone deacetylase; epigenetic regulation | Homo sapiens | 20Y | X-ray | 1.57 | Zn-dependent catalytic pocket |
| KMC11-05 | 3F3V | Src kinase | Non-receptor tyrosine kinase; cell signaling | Gallus gallus | 1BU | X-ray | 2.600 | c-Src kinase domain; S345C construct |
| KMC11-06 | 4YX4 | Carbonic anhydrase II | Zn metalloenzyme; CO<sub>2</sub>/bicarbonate conversion | Homo sapiens | FB2 | X-ray | 1.01 | Benzenesulfonamide inhibitor; catalytic Zn |
| KMC11-07 | 3PWH | Adenosine A2A receptor | GPCR; purinergic signaling | Homo sapiens | ZMA | X-ray | 3.296 | Thermostabilized receptor construct |
| KMC11-08 | 5FCK | Complement factor D | Serine protease; alternative complement pathway | Homo sapiens | 5WC | X-ray | 1.86 | Complement factor D inhibitor complex |
| KMC11-10 | 3LDP | GRP78/BiP | HSP70 chaperone; ER folding/stress response | Homo sapiens | 3P1 | X-ray | 2.20 | ATPase-domain inhibitor complex |
| KMC11-11 | 5LWM | JAK3 | Tyrosine kinase; cytokine/JAK-STAT signaling | Homo sapiens | 79T | X-ray | 1.55 | JAK3 kinase-inhibitor complex |

The benchmark covers a broad range of biological functions, including protein kinases (p38 MAPK, ABL1, Src, and JAK3), histone deacetylase 2 (HDAC2), carbonic anhydrase II, the adenosine A2A receptor, complement factor D, and the molecular chaperone GRP78/BiP. These proteins exhibit substantial structural diversity, encompassing different protein folds and ligand-binding environments, including kinase inhibitor pockets, zinc-dependent catalytic sites, a transmembrane GPCR binding pocket, and protease and chaperone binding sites. All ten holo structures were determined by X-ray crystallography, with resolutions ranging from 1.01 to 3.30 Å. Notably, the dataset includes wild-type ABL1 (3QRI) and its T315I gatekeeper mutant (3QRJ), both bound to the same inhibitor, DCC-2036, providing an opportunity to examine how a localized mutation affects molecular recognition.

The experimentally derived association rates span nearly four orders of magnitude, ranging from 2.50×10^2^ to 1.62×10^6^ M ¹s ¹ (Supplementary **Table S1**). The lowest association rate was associated with HDAC2 (4LY1), whereas the highest was associated with complement factor D (5FCK). Together, the diversity of protein families, ligand-binding environments, and association-rate scales provides a heterogeneous benchmark for evaluating the transferability of the multiscale Monte Carlo framework across distinct molecular systems. Specifically, the benchmark was designed to assess whether the simulations capture experimentally observed differences in association kinetics, distinguish the kinetic behaviors of encounter, pocket entry, and productive binding, and identify the physical determinants of molecular recognition through systematic interaction perturbations. Because the association rates were reconstructed from potentially nonidentical experimental conditions, the primary emphasis was placed on relative kinetic trends and mechanistic interpretation rather than precise prediction of absolute association rates.

### 3.2. Demonstrating protein–ligand association simulations by a representative case study

To illustrate the molecular recognition process captured by our multiscale Monte Carlo framework, we examined a representative association trajectory of the ABL1 kinase inhibitor DCC-2036 using the wild-type ABL1 structure [31] (PDB: 3QRI). As shown in **Figure 3a**, the ligand was initially positioned away from the protein surface and subsequently explored different spatial positions and orientations through stochastic translational, rotational, and torsional moves. During the 10,000 step trajectory, the ligand progressively approached the protein and established contact with the binding-pocket region. The structural snapshots illustrate the transition from an initially unbound configuration to a protein-associated state, demonstrating the ability of the simulation to capture ligand approach and pocket recognition.

**Figure 3.**
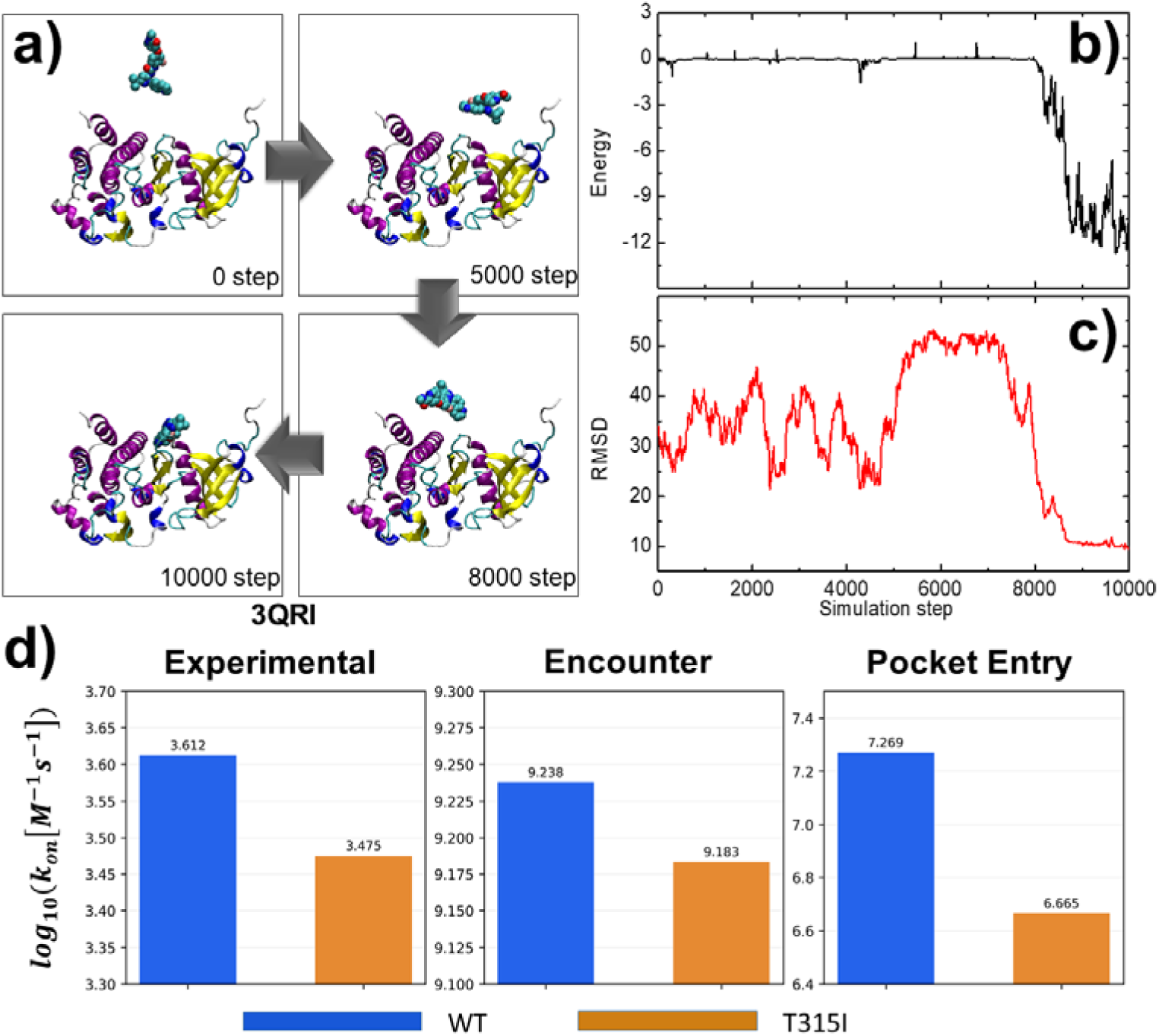
The representative simulation of protein–ligand association and comparison of wild-type and mutant ABL1 kinetics. (a) Structural snapshots from a representative Monte Carlo trajectory of DCC-2036 associating with wild-type ABL1 kinase (PDB: 3QRI), illustrating ligand configurations at 0, 5,000, 8,000, and 10,000 simulation steps. (b) Evolution of the effective protein-ligand interaction energy along the trajectory, showing the development of favorable interactions during ligand approach. (c) Corresponding ligand heavy-atom RMSD relative to the crystallographic bound conformation. The decrease in RMSD indicates progression toward the native binding region, although the productive-binding criterion was not reached within the displayed trajectory. (d) Comparison of experimentally derived association rates and simulated encounter and pocket-entry rates for wild-type ABL1 (3QRI) and its T315I mutant (3QRJ), both bound to DCC-2036. Rates are presented on a logarithmic scale in M ¹s ¹.

The corresponding interaction-energy profile remained close to zero during the early stages of the trajectory, consistent with the ligand exploring configurations with limited favorable protein contacts (**Figure 3b**). A pronounced decrease in energy occurred during the later stages, reaching approximately −10 to −12 kcal/mol as the ligand established more favorable interactions with the protein. The ligand RMSD relative to the experimental bound conformation exhibited substantial fluctuations during the initial search, followed by a marked decrease after approximately 8,000 simulation steps (**Figure 3c**). These changes indicate a transition from relatively unconstrained spatial exploration toward a more localized, energetically favorable protein-associated configuration. Although the ligand approached the native binding region, its RMSD remained approximately 10 Å at the end of the trajectory, indicating that the native binding pose had not been fully recovered within the observation window.

To further examine whether the framework can distinguish changes in molecular recognition arising from localized structural differences, we compared wild-type ABL1 (3QRI) with its T315I gatekeeper mutant (3QRJ), both bound to DCC-2036 (**Figure 3d**). The experimentally derived association rate decreased from (4.10×10^3^) to (2.99×10^3^) M ¹s ¹ upon mutation. Consistent with this direction of change, the simulated encounter rate decreased modestly, from (1.73×10^9^) to (1.53 ×10^9^) M ¹s ¹, whereas the pocket-entry rate decreased more substantially, from (1.86×10^7^) to (4.62×10^6^) M ¹s ¹. These results suggest that the T315I mutation has a greater effect on ligand access to the binding-pocket region than on initial protein–ligand encounter. Although the simulated pocket-entry difference exceeds the experimentally derived association-rate difference, the agreement in direction illustrates the sensitivity of the model to localized changes in the protein binding environment.

Together, these results demonstrate that the multiscale Monte Carlo framework can resolve successive stages of ligand approach and pocket recognition while providing stage-specific measures of association behavior. The representative trajectory and matched ABL1 comparison establish a basis for the subsequent benchmark-wide evaluation of association kinetics across structurally diverse protein–ligand systems.

### 3.3. Benchmark-wide evaluation of multistage protein–ligand association kinetics

To evaluate the kinetic behavior of the multiscale Monte Carlo framework across the benchmark, we compared the simulated encounter, pocket-entry, and productive-binding rates with experimentally derived association rate constants for all ten protein–ligand complexes. The three kinetic endpoints exhibited distinct relationships with experimental association rates, reflecting different levels of molecular recognition specificity. The simulated encounter rates were concentrated within a relatively narrow range, approximately 10^9^ M ¹s ¹, despite the nearly four-order-of-magnitude variation in experimental association rates (**Figure 4a**). Consequently, encounter rates exhibited relatively weak correlations with experiment, with a Pearson correlation coefficient of 0.475 and a Spearman rank correlation coefficient of 0.492 (**Table 2**). This limited discrimination suggests that initial protein–ligand contact alone does not adequately capture the kinetic differences among the benchmark complexes.

**Figure 4.**
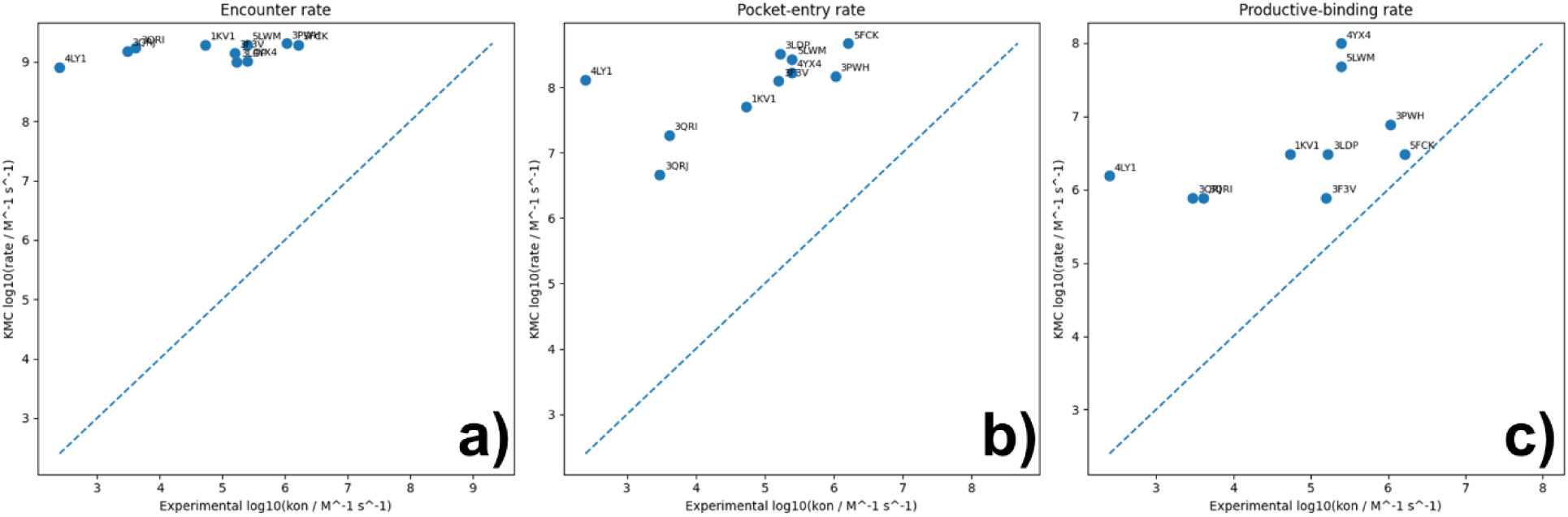
Benchmark-wide comparison of simulated association kinetics with experimentally derived association rates. Scatter plots comparing experimental association rate constants (*k_on_*) with simulated (a) encounter, (b) pocket-entry, and (c) productive-binding rates for the ten protein–ligand complexes. Both experimental and simulated rates are expressed as (*log_10_(k)*), with (k) in M ¹s ¹. Individual points represent protein-ligand complexes identified by their PDB codes. Dashed diagonal lines indicate equality between simulated and experimental rates. Pearson correlation coefficients, Spearman rank correlation coefficients, and prediction errors for each kinetic endpoint are summarized in Table 2. The experimentally derived association rates were reconstructed from independently reported dissociation rate constants and equilibrium dissociation constants.

**Table 2.** Comparison of simulated kinetic endpoints with experimental association rates. Statistical evaluation of encounter, pocket-entry, and productive-binding rates against experimentally derived association rates using Pearson and Spearman correlations, mean absolute error (MAE), and root-mean-square error (RMSE).

| Metric | Encounter | Pocket entry | Productive binding |
| --- | --- | --- | --- |
| Pearson r | 0.474842 | 0.616570 | 0.505616 |
| Spearman $\rho$ | 0.492403 | 0.757576 | 0.702080 |
| MAE ($\log_{10}$ units) | 4.402274 | 3.218870 | 1.822659 |
| RMSE ($\log_{10}$ units) | 4.538128 | 3.348501 | 2.082520 |

In contrast, pocket-entry rates displayed greater variation across protein–ligand systems and exhibited the strongest correlations with experimental association kinetics (Pearson correlation coefficient of 0.617 and Spearman rank correlation coefficient of 0.758 (**Figure 4b**)). This improvement suggests that ligand localization within the native binding-pocket region provides additional information about association behavior beyond nonspecific encounter. The greater dispersion of pocket-entry rates indicates that the structural and energetic requirements for accessing the binding pocket introduce system-dependent kinetic differences that are not fully captured by initial protein–ligand contact.

Introducing additional requirements for native-pose recovery and native-contact formation further reduced the apparent association rates. Productive-binding rates exhibited a Pearson correlation coefficient of 0.506 and a Spearman rank correlation coefficient of 0.702 with experimental *k_on_*(**Figure 4c**), indicating that substantial information about relative association behavior was retained despite the more stringent binding criterion. Importantly, the agreement in absolute rate magnitude improved progressively across the three kinetic endpoints. The mean absolute error decreased from 4.40 logarithmic units for encounter to 3.22 for pocket entry and 1.82 for productive binding, while the corresponding root-mean-square errors decreased from 4.54 to 3.35 and 2.08, respectively (**Table 2**). Thus, although pocket-entry rates provided the strongest correlation with experimental kinetics, productive-binding rates yielded the closest agreement in absolute magnitude.

Together, these results indicate that the different stages of protein–ligand association capture complementary aspects of molecular recognition. Initial encounter is relatively rapid across the benchmark and provides limited discrimination among systems, whereas pocket entry and productive native-like binding introduce additional structural and energetic constraints that distinguish their association behavior. Although the simulated rates generally overestimated experimental values, the positive correlations and progressive reduction in prediction errors support the utility of the multistage framework for characterizing relative association kinetics. Given the approximate simulation time mapping and uncertainties associated with experimentally reconstructed *k_on_*values, these comparisons are interpreted primarily as evidence of qualitative kinetic correspondence rather than quantitative validation of absolute association rates.

A recent multiscale study by Muñiz-Chicharro et al. (2025) combined Brownian dynamics (BD) and molecular dynamics (MD) to estimate protein–ligand association rates across ten complexes, reporting Spearman rank correlations of 0.24 for BD alone and 0.72 for the combined BD–MD approach [32]. Our pocket-entry rates exhibited a slightly higher Spearman correlation of 0.758, although these values cannot be directly compared because the two studies employed different benchmark systems, experimental measurements, and definitions of association. Notably, the previous benchmark comprised ten complexes involving four distinct protein targets, with multiple related ligands evaluated against the same target, whereas our benchmark includes ten complexes representing nine distinct protein targets across a broader range of protein families and binding-pocket environments. Furthermore, our experimentally derived association rates span approximately four orders of magnitude, compared with approximately two orders of magnitude in the previous study. Although the earlier benchmark benefits from directly measured association rates and controlled within-target ligand comparisons, our dataset provides broader coverage of biological targets and kinetic regimes. The comparable rank correlations, despite these differences in benchmark composition, suggest that our computationally simplified framework can capture meaningful relative association behavior across heterogeneous protein–ligand systems without requiring all-atom MD simulations.

### 3.4. Stage-specific physical determinants of protein–ligand association

To investigate the physical interactions governing different stages of protein–ligand association, we performed a systematic computational perturbation analysis across the ten benchmark complexes. Four molecular determinants were examined: electrostatic interactions, hydrophobic interactions, hydrogen bonding, and ligand conformational flexibility. For each determinant, three perturbation conditions were introduced relative to the baseline model: complete removal (0×), weakening (0.5×), and strengthening (2×), with the unperturbed condition (1×) serving as the reference. This resulted in 13 simulation conditions for each protein–ligand complex, comprising one baseline and 3×4=12 individual perturbations. Electrostatic, hydrophobic, and hydrogen-bond perturbations were implemented by scaling their respective contributions to the effective interaction energy, while keeping all other interaction terms unchanged. Ligand flexibility was independently perturbed by scaling the amplitude of internal torsional moves. All other simulation settings and kinetic-state definitions were maintained consistently across conditions.

For each perturbation condition, independent Monte Carlo trajectories were generated to evaluate changes in encounter, pocket-entry, and productive-binding kinetics. The perturbation response was quantified as the logarithmic rate difference, 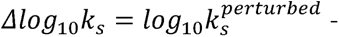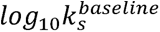, where (s) denotes one of the three association stages. Positive values indicate an increase in the apparent rate relative to baseline, whereas negative values indicate a decrease. The resulting perturbation profiles across all 12 conditions revealed distinct sensitivities across the three stages of molecular recognition.

Initial encounter kinetics exhibited relatively weak sensitivity to the perturbations examined (**Figure 5a**). Among most conditions, the average changes in encounter rates remained within approximately ±0.03 logarithmic units. Electrostatic and hydrophobic perturbations produced only minor changes, while increasing ligand flexibility resulted in a modest enhancement of encounter kinetics. These observations suggest that initial protein–ligand contact is primarily governed by diffusional exploration and geometric accessibility under the present simulation conditions, with relatively limited sensitivity to the short-range interaction terms included in the model. This finding is consistent with the narrow distribution of encounter rates observed across the benchmark in **Figure 4a**.

**Figure 5.**
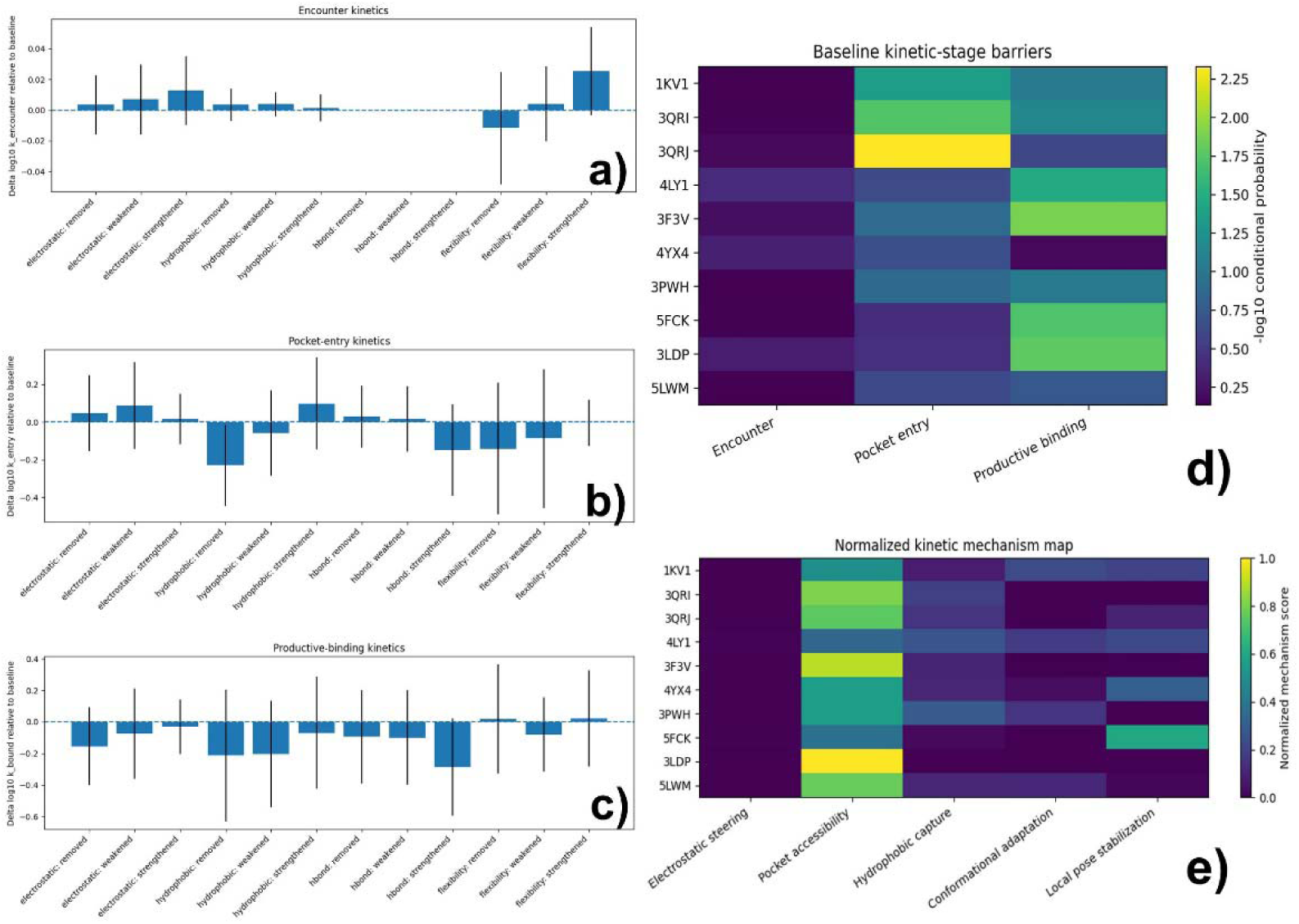
Sche Stage-specific physical determinants of protein–ligand association. Effects of perturbing electrostatic interactions, hydrophobic interactions, hydrogen bonding, and ligand flexibility on **(a)** encounter, **(b)** pocket-entry, and **(c)** productive-binding kinetics. Bars represent mean changes in ***log_10_*** association rates relative to baseline, with error bars indicating variability across complexes. Dashed lines indicate zero change. **(d)** Baseline kinetic-stage barriers calculated as the negative logarithm of conditional success probabilities, with higher values indicating greater difficulty in progressing through each stage. **(e)** Normalized kinetic mechanism map showing the relative contributions of five model-derived descriptors: electrostatic steering, pocket-entry difficulty, hydrophobic capture, conformational adaptation, and local pose stabilization.

In contrast, pocket-entry kinetics exhibited substantially greater sensitivity to the physical perturbations (**Figure 5b**). Removing hydrophobic interactions produced the largest average reduction in pocket-entry rates, with *Δlog*_10_*k_entry_* of approximately −0.23, corresponding to a decrease of about 41% relative to baseline. Weakening hydrophobic interactions also reduced pocket-entry rates, whereas strengthening them produced an average increase of approximately 0.10 logarithmic units. These trends suggest that hydrophobic attraction facilitates ligand localization within the native binding-pocket region. Ligand flexibility also influenced pocket entry: suppressing or weakening torsional motion decreased the apparent entry rates by approximately 0.14 and 0.09 logarithmic units, respectively. This observation is consistent with a role for internal conformational rearrangement in enabling ligands to access geometrically constrained binding environments. By comparison, electrostatic and hydrogen-bond perturbations produced relatively small average effects on pocket entry.

Productive-binding kinetics showed a different pattern of perturbation sensitivity (**Figure 5c**). Removing electrostatic interactions reduced the average productive-binding rate by approximately 0.16 logarithmic units, while removing or weakening hydrophobic interactions produced reductions of approximately 0.21 and 0.20 logarithmic units, respectively. These results suggest that favorable electrostatic and hydrophobic interactions contribute to the formation of native-like binding configurations after initial pocket recognition. Interestingly, strengthening hydrogen-bond interactions produced an even larger average reduction of approximately 0.28 logarithmic units, despite the expectation that stronger hydrogen bonding would favor complex stabilization. One possible reason is that excessive local attraction stabilizes nonproductive ligand orientations or intermediate configurations, thereby hindering progression toward the native binding pose. In contrast to pocket entry, perturbations of ligand flexibility produced relatively small average changes in productive-binding rates.

To further characterize differences in association behavior among individual protein–ligand systems, we examined the conditional probabilities of progressing through successive recognition stages using the unperturbed baseline simulations (**Figure 5d**). Three probability-derived kinetic-stage barriers were calculated as *B_s_* _=_ -*log*_10_*P_s_*, where *P_s_* denotes the conditional success probability for the corresponding stage. Specifically, the encounter barrier was derived from the fraction of trajectories reaching the encounter state, the pocket-entry barrier from the probability of pocket entry following encounter, and the productive-binding barrier from the probability of achieving native-like binding following pocket entry. Conditional probabilities were estimated using Jeffreys smoothing to account for finite trajectory counts and zero-event observations. Higher barrier values indicate lower conditional success probabilities and greater difficulty in progressing through the corresponding stage. The resulting heatmap revealed substantial heterogeneity among the ten complexes. The ABL1 complexes (3QRI and 3QRJ) exhibited relatively pronounced barriers during pocket entry, particularly for the T315I mutant, whereas Src kinase (3F3V), complement factor D (5FCK), and GRP78/BiP (3LDP) exhibited greater difficulty during productive-binding formation. In contrast, initial encounter generally showed lower probability-derived barriers across the benchmark. These findings suggest that the principal kinetic obstacles to molecular recognition vary among protein–ligand systems rather than being uniformly associated with a single stage.

Finally, to connect these stage-specific kinetic obstacles with their underlying physical determinants, we constructed a normalized kinetic mechanism map combining baseline conditional probabilities with the effects of removing individual interaction components (**Figure 5e**). Five operational descriptors were defined: electrostatic steering, pocket-entry difficulty, hydrophobic capture, conformational adaptation, and local pose stabilization. Electrostatic steering was quantified as the positive logarithmic reduction in encounter probability following removal of electrostatic interactions, whereas pocket-entry difficulty was defined as the negative logarithm of the baseline conditional pocket-entry probability. Hydrophobic capture was quantified from the reduction in conditional pocket-entry probability upon removal of hydrophobic attraction. Conformational adaptation and local pose stabilization were similarly evaluated from reductions in conditional productive-binding probability following suppression of ligand torsional motion and removal of hydrogen-bond interactions, respectively. For each complex, the five nonnegative scores were normalized by their sum to obtain a relative mechanistic profile.

The resulting map revealed heterogeneous recognition profiles across the benchmark. For example, the ABL1 complexes (3QRI and 3QRJ) exhibited prominent pocket-entry difficulty scores, consistent with the relatively low conditional pocket-entry probabilities identified in Fig. 5d. In contrast, complement factor D (5FCK) showed a comparatively prominent local pose stabilization score, suggesting that productive-binding success in this system is particularly sensitive to the removal of hydrogen-bond interactions. Several other complexes, including p38 MAP kinase (1KV1) and the adenosine A2A receptor (3PWH), exhibited detectable hydrophobic-capture contributions, indicating that hydrophobic attraction may facilitate ligand progression into their binding regions. Electrostatic steering and conformational adaptation generally contributed smaller normalized scores across the benchmark under the present simulation conditions. These examples illustrate how different protein–ligand systems can exhibit distinct combinations of kinetic obstacles and sensitivities to individual physical interactions. Because the descriptors combine baseline kinetic difficulty with perturbation-derived sensitivities, their normalized values represent operational, model-dependent measures rather than directly comparable physical energy contributions.

Taken together, the computational perturbation results reveal a stage-dependent organization of the physical determinants of protein–ligand recognition. Initial encounter was comparatively insensitive to the perturbations examined, whereas pocket entry showed greater dependence on hydrophobic attraction and ligand conformational flexibility. Productive binding exhibited additional sensitivity to electrostatic, hydrophobic, and hydrogen-bond interactions, including potentially unfavorable effects of excessive local stabilization. The substantial variation among individual complexes further suggests that different molecular systems encounter distinct obstacles along their association pathways. Although the relatively large variability in several perturbation responses limits definitive assignment of individual physical mechanisms, these results demonstrate how systematic computational perturbations can be used to construct an interpretable, stage-resolved description of association between proteins and small-molecular ligands.

## 4. Concluding Discussions

Protein–small-molecule association is a fundamental molecular recognition process underlying numerous biological functions and therapeutic interventions. The kinetics of this process influence how rapidly ligands engage their targets, thereby affecting drug efficacy, selectivity, and pharmacological responses. Understanding the physical mechanisms governing association is therefore essential for elucidating molecular recognition and guiding rational drug design. However, achieving this understanding requires moving beyond static binding structures and equilibrium affinities to characterize the dynamic pathways and intermolecular interactions through which ligands recognize and engage their protein targets. In order to reach this goal, here we developed a multiscale Monte Carlo framework that combines a flexible, atomistic ligand with a coarse-grained protein binding pocket and an atomistic steric envelope. This hybrid representation enables efficient sampling of ligand translational, rotational, and internal conformational motions while retaining an interpretable description of the physical interactions governing molecular recognition. By resolving association into encounter, pocket entry, and productive binding, the framework provides a stage-specific perspective on protein–ligand recognition that complements conventional approaches focused on a single overall association rate.

Evaluation across ten protein–ligand complexes representing nine distinct protein targets revealed positive correlations between simulated association behavior and experimentally derived association rates. Among the three kinetic endpoints, pocket-entry rates exhibited the strongest correspondence with experiment, with a Pearson correlation coefficient of 0.617 and a Spearman rank correlation coefficient of 0.758. In contrast, initial encounter rates varied relatively little across the benchmark, suggesting that the kinetic differences among these systems emerge primarily after initial protein–ligand contact. Productive-binding rates exhibited smaller deviations from experimental rate magnitudes, although their rank correlations were somewhat weaker than those of pocket entry. These observations suggest that the structural and energetic requirements associated with accessing the native binding region provide important information about the diversity of protein–ligand association behavior. Nevertheless, the remaining discrepancies in absolute rates indicate that the present framework should be viewed primarily as a tool for comparative kinetic analysis rather than a quantitatively calibrated predictor of association rate constants.

A central contribution of this work is the use of systematic computational perturbations to investigate the physical determinants of individual recognition stages. By independently removing, weakening, or strengthening electrostatic interactions, hydrophobic attraction, hydrogen bonding, and ligand torsional flexibility, we observed distinct kinetic responses across the association pathway. Initial encounter was comparatively insensitive to these perturbations, whereas pocket entry showed greater dependence on hydrophobic attraction and ligand conformational flexibility. Productive binding exhibited additional sensitivity to electrostatic and local interactions. Interestingly, strengthening hydrogen bonding did not uniformly accelerate productive binding and instead reduced its average apparent rate. This observation raises the possibility that excessively favorable local interactions may stabilize nonproductive configurations and impede progression toward the native binding pose. Although direct characterization of such kinetic traps remains necessary, these findings illustrate that strengthening favorable molecular interactions does not necessarily accelerate association.

Several limitations of the present framework should be acknowledged. First, the protein was treated as structurally rigid, preventing explicit representation of conformational changes, induced fit, and binding-site gating during molecular recognition. Future improvements could incorporate protein flexibility through an elastic network model, allowing collective protein motions and local binding-pocket rearrangements while preserving the computational efficiency of the coarse-grained representation. Second, solvent effects were represented only approximately through screened electrostatic interactions and an effective hydrophobic potential, without explicitly accounting for solvent reorganization, hydration-shell dynamics, or water-mediated interactions during ligand association. This limitation could be addressed by introducing an implicit-solvent description that accounts for electrostatic and nonpolar solvation contributions, supplemented by localized hydration-dependent potentials to capture the energetic consequences of water displacement and rearrangement within the binding pocket.

Beyond providing mechanistic insights into protein–ligand recognition, the framework offers several potential applications in computational drug discovery and molecular design. By distinguishing the physical interactions and kinetic stages that govern ligand association, it could help identify molecular features that facilitate rapid target engagement or hinder productive binding, providing complementary guidance to conventional affinity-based drug optimization. The computational efficiency and modularity of the approach also make it suitable for integration with machine learning and optimization algorithms, enabling the development of physics-informed strategies for exploring ligand modifications and their effects on association pathways. Furthermore, the framework could be extended to more complex molecular recognition processes, particularly molecular-glue-induced ternary complex formation, where sequential and cooperative binding events play important roles in therapeutic activity. Ultimately, connecting molecular interaction mechanisms with stage-specific association behavior may provide a foundation for designing therapeutic molecules with optimized binding kinetics, selectivity, and functional responses.

## Supporting information

Kinetic provenance and structural mapping of the simulation-ready benchmark

## Acknowledgment

This work was supported by the National Institutes of Health under grant number R01GM162824, the United States–Israel Binational Science Foundation Project Number: 2023336, and the Einstein 2030 Seed Fund. The work was also partially supported by a start-up grant from Albert Einstein College of Medicine. Computational support was provided by Albert Einstein College of Medicine High Performance Computing Center.

## Author Contributions

Z.S. and Y.W. designed research; Z.S. and Y.W. performed research; Y.W. analyzed data; Y.W. wrote the paper.

## Competing financial interests

The authors declare no competing financial interests.

## Notes

### Competing Interest Statement

The authors have declared no competing interest.

