## Supplementary material for "Decoding the Physical Mechanisms of Protein-Small Molecule Association by Multiscale Kinetic Monte-Carlo Simulations": Kinetic provenance and structural mapping of the simulation-ready benchmark

**Supplementary Table S1**. Kinetic provenance and structural mapping of the simulation-ready benchmark

*Experimental koff values were obtained from the Liu et al. curated dissociation-kinetics dataset; KD values were matched from BindingDB. Derived kon was calculated as koff / KD(M). Because koff and KD can originate from different experiments and conditions, derived kon values are treated as approximate kinetic references rather than condition-matched direct measurements.*

| **ID** | **PDB** | **UniProt** | **Target** | **Ligand CCD** | **koff (s^-1)** | **koff method** | **Temp. (°C)** | **KD median (nM)** | **KD range (nM)** | **Derived kon (M^-1 s^-1)** | **log10 kon** | **No. KD records** | **Kinetic match** | **PDB ligand match** | **BindingDB row(s)** | **BindingDB DOI(s)** | **koff reference** |
| --- | --- | --- | --- | --- | --- | --- | --- | --- | --- | --- | --- | --- | --- | --- | --- | --- | --- |
| KMC11_01 | 1KV1 | Q16539 | P38α | BMU | 0.062 | Surface plasmon resonance | NR | 1160 | 1160 | 5.34e+04 | 4.728 | 3 | exact inchikey+uniprot | Exact | 27477; 27480; 27489 | 10.1021/jm030120s; 10.1038/nsb770 | Biochemistry. 2010, 49, 3611-3618 |
| KMC11_02 | 3QRI | P00519 | abl1 | 919 | 2.95e-05 | Slow-binding inhibition | NR | 7.2 | 7.2 | 4097.22 | 3.612 | 1 | exact inchikey+uniprot | Exact | 73887 | 10.1021/acschembio.5b01018 | Cancer Cell. 2011, 19, 556–568 |
| KMC11_03 | 3QRJ | P00519 | abl1 | 919 | 2.15e-05 | Slow binding inhibition | NR | 7.2 | 7.2 | 2986.11 | 3.475 | 1 | exact inchikey+uniprot | Exact | 73887 | 10.1021/acschembio.5b01018 | Cancer Cell. 2011, 19, 556–568 |
| KMC11_04 | 4LY1 | Q92769 | histone deacetylase 2 | 20Y | 1.38e-05 | Proteros reporter displacement | NR | 55.2 | 55.2 | 250 | 2.398 | 1 | exact inchikey+uniprot | Exact | 76821 | 10.1074/jbc.M113.490706 | J. Biol. Chem. 2013, 288:26926-26943 |
| KMC11_05 | 3F3V | P00523 | Proto-oncogene tyrosine-protein kinase Src | 1BU | 0.0115 | Fluorescent-labeled kinase assay | NR | 73 | 73 | 1.58e+05 | 5.197 | 1 | exact inchikey+uniprot | Exact | 56338 | 10.1021/jm9002928 | J. Med. Chem. 2009, 52, 3915–3926 |
| KMC11_06 | 4YX4 | P00918 | Human Carbonic Anhydrase II | FB2 | 0.163 | Surface plasmon resonance | NR | 660 | 660 | 2.47e+05 | 5.393 | 1 | exact inchikey+uniprot | Exact | 22691 | 10.1021/ja057257n | J Med Chem. 2016, 59, 4245-4256 |
| KMC11_07 | 3PWH | P29274 | Thermostabilised Adenosine A2A Receptor | ZMA | 0.000233 | Radioligand displacement assay | 4 | 0.22 | 0.22 | 1.06e+06 | 6.025 | 2 | exact inchikey+uniprot | Exact | 38489; 38512 | 10.1021/jm701185v; 10.1021/jm701187w | J. Med. Chem. 2016, 59, 6470−6479 |
| KMC11_08 | 5FCK | P00746 | Complement factor D | 5WC | 0.0102 | Surface plasmon resonance | 25 | 6.29 | 6.03-6.55 | 1.62e+06 | 6.210 | 2 | connectivity+uniprot | Exact | 75996; 75997 | 10.1038/nchembio.2208 | Nat Chem Biol. 2016, 12, 1105-1110 |
| KMC11_10 | 3LDP | P11021 | Glucose Regulated Protein | 3P1 | 0.4 | Surface plasmon resonance | 25 | 2410 | 2410 | 1.66e+05 | 5.220 | 1 | exact inchikey+uniprot | Connectivity | 62971 | 10.1021/jm101625x | J. Med. Chem. 2011, 54, 4034–4041 |
| KMC11_11 | 5LWM | P52333 | Tyrosine-protein kinase JAK3 | 79T | 0.00033 | Proteros reporter displacement | NR | 1.34 | 1.34 | 2.46e+05 | 5.391 | 1 | connectivity+uniprot | Connectivity | 75760 | 10.1016/j.chembiol.2016.10.008 | Cell Chem Biol. 2016, 23, 1335-1340 |

**Notes.** NR, not reported in the curated source table. “Kinetic match” describes the ligand/target matching used to connect the Liu et al. koff record to BindingDB; “PDB ligand match” describes the ligand identity match used for the simulation-ready holo structure. The 5A3O holo candidate was not included in the KMC benchmark because its experimental ligand is represented by multiple PDB chemical components.

| **Column** | **Description** |
| --- | --- |
| **ID** | Internal benchmark identifier assigned to each simulation-ready protein–ligand complex. |
| **PDB** | Four-character Protein Data Bank identifier of the experimentally determined holo structure used for simulation. |
| **UniProt** | UniProt accession identifying the protein target used to match kinetic and affinity records. |
| **Target** | Protein name or construct associated with the benchmark complex. |
| **Ligand CCD** | PDB Chemical Component Dictionary code identifying the bound small-molecule ligand. |
| **koff (s⁻¹)** | Experimentally reported ligand dissociation rate constant obtained from the curated dissociation-kinetics dataset. |
| **koff method** | Experimental assay or technique used to determine the reported dissociation rate. |
| **Temp. (°C)** | Temperature reported for the dissociation-kinetics measurement; NR means not reported in the curated source table. |
| **KD median (nM)** | Median of matched BindingDB equilibrium dissociation constants, expressed in nanomolar units. |
| **KD range (nM)** | Minimum-to-maximum range of matched BindingDB KD records; a single value indicates no range across the retained records. |
| **Derived kon (M⁻¹ s⁻¹)** | Approximate association rate calculated as koff / KD, with KD converted from nM to molar units. |
| **log10 kon** | Base-10 logarithm of the derived association rate in M⁻¹ s⁻¹. |
| **No. KD records** | Number of matched BindingDB KD measurements used to summarize affinity for the protein–ligand pair. |
| **Kinetic match** | Identity-matching rule connecting the curated koff record to BindingDB, using ligand InChIKey and protein UniProt identity; connectivity-only matches do not verify stereochemistry. |
| **PDB ligand match** | Level of chemical-identity agreement between the kinetic ligand and the ligand in the holo PDB structure: Exact or Connectivity (matching connectivity without full stereochemical confirmation). |
| **BindingDB row(s)** | BindingDB record identifiers for the KD entries included in the affinity matching. |
| **BindingDB DOI(s)** | Digital object identifier(s) of the publication(s) associated with the matched BindingDB affinity measurements. |
| **koff reference** | Literature source of the experimental dissociation-rate measurement. |
